# Exploratory Correlation of The Human Structural Connectome with Non-MRI Variables in Alzheimer’s Disease

**DOI:** 10.1101/2023.06.30.547308

**Authors:** Iman Aganj, Jocelyn Mora, Aina Frau-Pascual, Bruce Fischl, the Alzheimer’s Disease Neuroimaging Initiative

## Abstract

**INTRODUCTION:** Discovery of the associations between brain structural connectivity and clinical and demographic variables can help to better understand the vulnerability and resilience of the brain architecture to neurodegenerative diseases and to discover biomarkers.

**METHODS:** We used four diffusion-MRI databases, three related to Alzheimer’s disease, to exploratorily correlate structural connections between 85 brain regions with non-MRI variables, while stringently correcting the significance values for multiple testing and ruling out spurious correlations via careful visual inspection. We repeated the analysis with brain connectivity augmented with multi-synaptic neural pathways.

**RESULTS:** We found 85 and 101 significant relationships with direct and augmented connectivity, respectively, which were generally stronger for the latter. Age was consistently linked to decreased connectivity, and healthier clinical scores were generally linked to increased connectivity.

**DISCUSSION:** Our findings help to elucidate which structural brain networks are affected in Alzheimer’s disease and aging and highlight the importance of including indirect connections.

## 1. Introduction

Normal aging, as well as debilitating neurodegenerative diseases such as Alzheimer’s disease (AD), affect not only individual brain regions, but also connectivity between them [1, 2]. Focus on brain regions, but not interregional connectivity, may have hindered progress in understanding and treating diseases such as AD that are characterized as disconnection syndromes [3]. Mapping the complex brain networks through which information flows – i.e., the human *connectome* [4] – can help to better understand the vulnerability and resilience of these networks to the effects of AD, potentially leading to the discovery of diagnostically and therapeutically important connectomic biomarkers. Analysis of structural brain networks, by means of noninvasive diffusion-weighted magnetic resonance imaging (dMRI), has proved valuable in revealing the structural basis of dysfunction in mild cognitive impairment (MCI) and AD, demonstrating changes distinct from those with healthy aging [5-9].

Brain connectivity is often represented as a graph adjacency matrix of connection strengths between the brain regions of interest (ROIs), with its number of elements (graph edges) growing quadratically with respect to the number of ROIs (graph nodes). In population connectomic studies, it is often desired to find links between brain connectivity and non-MRI (clinical and/or demographic) variables. Such a study typically has sufficient statistical power to test pre-hypothesized relationships involving specific brain connections and variables. In contrast, an exploratory investigation to *discover* previously unknown relationships would require correlating the connectivity strength of every brain ROI pair with every available variable, amounting to hundreds of thousands (sometimes millions) of tests. In that scenario, the correction for multiple comparisons would make the study statistically less powerful and consequently less desirable to conduct. Alternatively, one could reduce the number of tests considerably by focusing on network summary features [10] rather than brain connections, which would inform about how the variables relate to the network as a whole [7, 11] but not to individual brain connections.

Structural connectivity between two brain regions is commonly defined based on the dMRI tractography-derived [12, 13] streamlines between them. The direct fiber bundle connecting two brain areas is expected to be the major signal carrier between them; however, *multi-synaptic* neural pathways (those mediated through other regions) also provide connectivity [14, 15]. We have previously developed computational methods to augment direct structural connectivity graphs with indirect connections [16] as well as quantify brain structural connectivity while accounting for indirect pathways [17], and have shown the importance of these pathways in predicting functional connectivity [17] and deriving connectomic biomarkers for MCI and AD [18].

Here, we take an exploratory approach to discovering relationships that individual structural connections in the brain may have with clinical and demographic variables. We use anatomical and diffusion MR images along with non-MRI data from four public databases (three of which are related to AD) to find links between brain connections – both direct and augmented – and non-MRI variables that remain significant after stringent correction for multiple testing and visual inspection.

We describe our processing and analysis methods in Section 2, report our results in Section 3, discuss them in Section 4, and conclude the paper in Section 5.

## 2. Methods

### 2.1. Datasets

We used the following four public dMRI databases. The number of subjects indicates the subset of subjects that were processed and included in our analysis, and the number of non-MRI variables indicates variables that were available for at least some of the included subjects.

- The second phase of the *Alzheimer’s Disease Neuroimaging Initiative (****ADNI-2***) [19]: 217 subjects (from cognitively normal to AD), 47 non-MRI variables from the ADNIMERGE table (demographics, CSF markers, dementia/cognitive exam scores, PET, ApoE4, diagnosis,…).
- The third release in the *Open Access Series of Imaging Studies (****OASIS-3***) [20]: 771 subjects (from cognitively normal to AD), 588 non-MRI variables (demographics, Uniform Data Set, dementia/cognitive exam scores, ApoE,…).
- The *Pre-symptomatic Evaluation of Experimental or Novel Treatments for Alzheimer’s Disease (****PREVENT-AD***) [21]: 340 cognitively unimpaired older individuals with a parental or multiple-sibling history of AD, 199 non-MRI variables (demographics, medical history, vitals, CSF markers, dementia/cognitive exam sub-scores, genetics, lab results, auditory/olfactory processing,…).
- The WashU-UMN *Human Connectome Project (****HCP***) [22]: 617 healthy young adults, 488 non-MRI variables (demographics, medical history, family history, dementia/cognitive exam scores, personality/emotion tests, motor/sensory tests, task performance,…).

### 2.2. Data processing

Anatomical MR images of the databases were processed with FreeSurfer [23]. All time points of PREVENT-AD were also more robustly processed using the FreeSurfer longitudinal pipeline [24]. For all databases, we included each subject only once, i.e. the earliest visit containing dMRI (frequently the baseline), in order to keep our analyzed data points independent and our study cross-sectional. We then ran the FreeSurfer dMRI processing pipeline, which also includes commands from the FMRIB Software Library (FSL) [25], and propagated 85 automatically segmented cortical and subcortical regions from the structural to the diffusion space using boundary-based image registration [26].

Next, we used our public toolbox (www.nitrc.org/projects/csaodf-hough) to: 1) reconstruct the diffusion orientation distribution function in constant solid angle [27], 2) run Hough-transform global probabilistic tractography [13] to generate an optimal (highest-score) streamline passing through each of the 10,000 seed points per subject, 3) compute a symmetric structural connectivity matrix (with positive elements) for each subject by summing the tracts passing through each pair of ROIs weighted by the tract score (hence an emphasis on streamlines best aligned with the dMRI-derived fiber orientations as well as fiber tracts with the highest white-matter integrity), and 4) augment the raw matrices with indirect connections (see Section 2.3) [16]. We transformed the connectivity value *c* (each element in the raw or augmented connectivity matrix) as 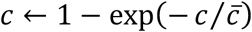, where 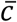 is the cross-subject average of *c*, thereby confining the connectivity values to the range [0,1).

### 2.3. Augmentation of structural connectivity with indirect connections

Strong functional connectivity between brain regions are commonly observed between regions with no *direct* structural connection [14, 28-34]. Some variance in functional connectivity unexplained by direct connections can be accounted for by *indirect* structural connections [14, 15, 17], implying that the network nature of the brain makes the interaction between two brain areas sensitive to influences from other remote areas [29].

We have previously developed a method to augment a tractography-generated structural connectivity matrix with indirect connections via the mathematics of circuits laws [16], thereby producing a new matrix that additionally reflects multi-synaptic pathways. This approach is based on the intuition that total connectivity for multiple direct connections is expectedly their sum if they are parallel, or smaller than each connection if they are in series (as total connectivity is presumably bottlenecked by the weakest link along the way). These conditions are accommodated by modeling the brain similarly to a resistive electrical circuit, where a resistor represents each direct connection, with its conductance (inverse of resistance) being the tractography-measured strength of the connection [16, 35]. Total (augmented) connectivity is then calculated via Kirchhoff’s laws as the overall conductance among regions, using graph Laplacian methods.

### 2.4. Analysis

We used the cross-sectional data of each database to independently test if there is a statistically significant relationship between each non-MRI (clinical or demographic) variable and the computed structural brain connection between each ROI pair. To perform a true exploratory analysis, we did not exclude any available variable based on its perceived relevance. To deal with data source heterogeneity, we analyzed the databases and report their results separately. The homogeneity within each database is expected to lead to findings that would be strengthened if they independently replicated in several databases.

If a non-MRI variable had categorical (rather than numeric) values, we converted it to numeric by assigning a natural number to each category, while making our best effort to sort the categories (if more than two) in a monotonic order; for instance, for *Baseline Diagnosis* in ADNI-2, we assigned: Control Normal → 1, Significant Memory Concern → 2, Early MCI → 3, Late MCI → 4, and AD → 5. We computed the Pearson correlation coefficient (*r*), along with its significance (*p*) value, between each variable and each connection. The *p*-values were then corrected for multiple comparisons via the conservative Bonferroni method (*p*_*b*_); i.e., they were multiplied by the number of (undirected) connections, #ROIs×(#ROIs-1)÷2 = 85×84÷2 = 3570, as well as by the number of studied variables (see Section 2.1). Since the quantified structural connectivity, which is the score-weighted number of streamlines passing through a pair of ROIs, is affected by the tract length, we controlled for the extraneous variable of intracranial volume (ICV) by computing the *partial* correlation instead. For robustness of the correlation [36], we removed connectivity values that were marginal outliers from the correlation analysis by excluding any element in the connectivity matrix of a subject (but not the subject’s entire matrix) that was larger than 0.9 (recall the range [0,1) of values). This was principally because we deemed a connectivity value with a large deviation from the population mean less reliable, since such a deviation indicated an increased likelihood of lower image quality or a data-processing issue. Therefore, slightly different numbers of subjects contributed to the correlation analysis of different brain connections.

For each variable, we selected the connection most significantly correlating with it, i.e. with the lowest *p*_*b*_-value. If *p*_*b*_ was smaller than the threshold α=0.05, then we scatter-plotted the connection strength with respect to the variable and visually inspected it to ensure the significant Pearson correlation was real and not spurious due to some outliers, thus avoiding situations with most data points clustered together with no obvious relationship [36]. The correlations surviving the Bonferroni correction and passing the visual inspection are reported as follows.

## 3. Results

Cross-subject medians of the raw and augmented connectivity matrices are shown in Figure 1 for the four databases. We correlated 3570 brain structural connections with 47, 588, 199, and 488 non-MRI variables for each of the ADNI-2, OASIS-3, PREVENT-AD, and HCP databases, respectively, while controlling for the ICV. Out of those variables, 15, 230, 32, and 84, respectively, were found to have significant Pearson correlation (*p*_*b*_ < 0.05) with raw connections, and 20, 79, 1, and 1 variables, respectively, had significant correlation with augmented connections. After visual inspection to remove spurious correlations, variables with significant correlation with raw connectivity were reduced to 15, 65, 3, and 2, respectively, whereas the variables significantly correlated with augmented connectivity remained unchanged. The findings are detailed in Tables 1 through 4 for the four databases. The right column indicates the total number (out of 3570) of connections reaching Bonferroni-level significance for a given variable.

**Table 1:** Significant correlations of non-MRI variables with brain connectivity in ADNI-2.

| Non-MRI variable | Most correlated brain structural connection | # Sig. Conn |
| --- | --- | --- |
| Baseline Diagnosis | L. Lingual cortex – L. Entorhinal cortex<br>*Augmented, $r = -0.41, p_b = 0.0005$* | 79 |
| Age | L. Hippocampus – L. Ventral diencephalon<br>*Raw, $r = -0.42, p_b = 0.0002$* | 7 |
|  | L. Hippocampus – R. Superior frontal cortex<br>*Augmented, $r = -0.47, p_b = 7 \times 10^{-7}$* | 67 |
| Sex | R. Putamen – Brainstem<br>*Raw, $r = 0.36$ (with the male sex), $p_b = 0.04$* | 2 |
| FDG-PET<br>*Mean of angular, temporal, and posterior cingulate* | R. Hippocampus – R. Fusiform cortex<br>*Raw, $r = 0.43, p_b = 0.0001$* | 3 |
|  | R. Hippocampus – R. Precuneus cortex<br>*Augmented, $r = 0.46, p_b = 4 \times 10^{-6}$* | 67 |
| AV45 PET (binding to β-amyloid)<br>*Mean of whole cerebellum* | R. Hippocampus – R. Precuneus cortex<br>*Augmented, $r = -0.43, p_b = 8 \times 10^{-5}$* | 45 |
| Clinical Dementia Rating (CDR)<br>*Sum of boxes* | R. Hippocampus – R. Fusiform cortex<br>*Raw, $r = -0.38, p_b = 0.007$* | 3 |
|  | R. Entorhinal cortex – L. Pallidum<br>*Augmented, $r = -0.43, p_b = 6 \times 10^{-5}$* | 135 |
| AD Assessment Scale (ADAS)<br>*11 items* | R. Hippocampus – R. Fusiform cortex<br>*Raw, $r = -0.37, p_b = 0.03$* | 2 |
|  | R. Hippocampus – R. Precuneus cortex<br>*Augmented, $r = -0.43, p_b = 0.0001$* | 45 |
| AD Assessment Scale (ADAS)<br>*13 items* | R. Hippocampus – R. Fusiform cortex<br>*Raw, $r = -0.39, p_b = 0.005$* | 2 |
|  | R. Hippocampus – R. Precuneus cortex<br>*Augmented, $r = -0.44, p_b = 4 \times 10^{-5}$* | 86 |
| AD Assessment Scale (ADAS)<br>*Delayed Word Recall* | R. Hippocampus – R. Ventral diencephalon<br>*Raw, $r = -0.38, p_b = 0.01$* | 4 |
|  | L. Entorhinal cortex – R. Caudal anterior cingulate cortex<br>*Augmented, $r = -0.40, p_b = 0.0008$* | 42 |
| Mini-Mental State Examination (MMSE) | R. Hippocampus – R. Entorhinal cortex<br>*Raw, $r = 0.37, p_b = 0.02$* | 2 |
|  | L. Amygdala – R. Entorhinal cortex<br>*Augmented, $r = 0.42, p_b = 0.0003$* | 75 |
| Rey Auditory Verbal Learning Test (RAVLT) Immediate<br>*Sum of 5 trials* | R. Hippocampus – R. Entorhinal cortex<br>*Raw, $r = 0.39, p_b = 0.006$* | 1 |
|  | R. Isthmus cingulate cortex – L. Entorhinal cortex<br>*Augmented, $r = 0.41, p_b = 0.0005$* | 70 |
| Functional Assessment Questionnaire (FAQ) | R. Hippocampus – R. Fusiform cortex<br>*Raw, $r = -0.38, p_b = 0.01$* | 2 |
|  | R. Hippocampus – R. Rostral middle frontal cortex<br>*Augmented, $r = -0.46, p_b = 10^{-6}$* | 142 |
| Montreal Cognitive Assessment (MoCA) | L. Hippocampus – L. Middle temporal cortex<br>*Raw, $r = 0.39, p_b = 0.01$* | 2 |
|  | R. Hippocampus – R. Isthmus cingulate cortex<br>*Augmented, $r = 0.44, p_b = 6 \times 10^{-5}$* | 71 |
| ADNI modified Preclinical Alzheimer's Cognitive Composite (PACC)<br>*with Digit Symbol Substitution* | R. Hippocampus – R. Entorhinal cortex<br>*Raw, $r = 0.40, p_b = 0.001$* | 6 |
|  | R. Hippocampus – L. Rostral middle frontal cortex<br>*Augmented, $r = 0.44, p_b = 2 \times 10^{-5}$* | 161 |
| ADNI modified Preclinical Alzheimer's Cognitive Composite (PACC)<br>*with Trails B* | R. Parahippocampal cortex – R. Fusiform cortex<br>*Raw, $r = 0.41, p_b = 0.0005$* | 6 |
|  | R. Hippocampus – L. Rostral middle frontal cortex<br>*Augmented, $r = 0.44, p_b = 10^{-5}$* | 173 |
| Everyday Cognition Study Partner Report (ECog SP) – *Memory* | L. Entorhinal cortex – L. Banks of superior temporal sulcus<br>*Augmented, $r = -0.37, p_b = 0.02$* | 1 |
| Everyday Cognition Study Partner Report (ECog SP) – *Language* | L. Isthmus cingulate cortex – L. Middle temporal cortex<br>*Augmented, $r = -0.38, p_b = 0.01$* | 4 |
| Everyday Cognition Study Partner Report (ECog SP) – *Plan* | R. Isthmus cingulate cortex – L. Inferior temporal cortex<br>*Augmented, $r = -0.39, p_b = 0.009$* | 2 |
| Everyday Cognition Study Partner Report (ECog SP) – *Total* | L. Hippocampus – L. Entorhinal cortex<br>*Raw, $r = -0.37, p_b = 0.04$* | 1 |
|  | R. Isthmus cingulate cortex – L. Inferior temporal cortex<br>*Augmented, $r = -0.39, p_b = 0.006$* | 4 |
| Logical Memory<br>*Delayed Recall* | R. Hippocampus – L. Rostral middle frontal cortex<br>*Augmented, $r = 0.37, p_b = 0.01$* | 6 |
| Trail Making Test, Part B<br>*Time to complete* | R. Parahippocampal cortex – R. Fusiform cortex<br>*Raw, $r = -0.40, p_b = 0.001$* | 1 |
|  | R. Hippocampus – R. Isthmus cingulate cortex<br>*Augmented, $r = -0.37, p_b = 0.04$* | 1 |

**Figure 1.**
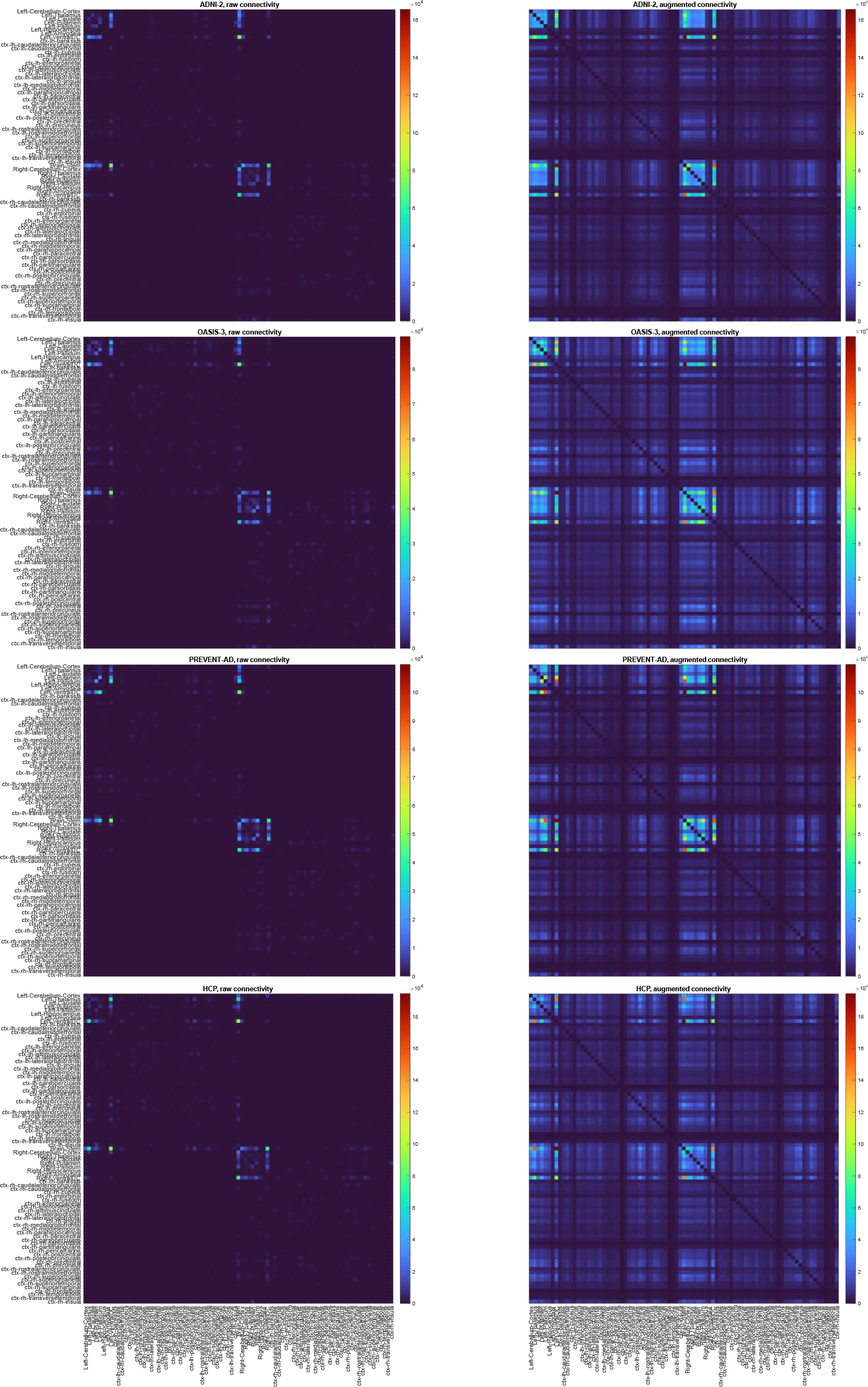
Element-wise cross-subject median of the raw (left) and augmented (right) connectivity matrices for the four databases.

Controlling for ICV had several effects on the results, e.g., it made the correlation of brain connectivity with *ECog SP – Memory* in ADNI-2 and with *Multilingual Naming Test (MINT*, both *the total score* and *the total correct without semantic cue)* in OASIS-3 significant (see Table 1 and Table 2). Without separating the effects of ICV, conversely, we would observe significant correlations of brain connectivity with *grip strength* (-) and the *maximum number of drinks consumed in a single day* (-) in HCP, and with *posture issues* (-) and *MoCA: Abstraction* (+) in OASIS-3. The confounding effect of ICV was especially drastic on the correlation with sex. Significant correlation of connectivity (of the most related brain connection) with the *male sex* was:

**Table 2:** Significant correlations of non-MRI variables with brain connectivity in OASIS-3.

| Non-MRI variable | Most correlated brain structural connection | # Sig. Conn |
| --- | --- | --- |
| Age | L. Hippocampus – L. Thalamus<br>*Raw*, $r = -0.48$, $p_b = 9 \times 10^{-35}$ | 266 |
| | L. Hippocampus – R. Lingual cortex<br>*Augmented*, $r = -0.5$, $p_b = 3 \times 10^{-41}$ | 2232 |
| Sex | R. Thalamus – L. Thalamus<br>*Raw*, $r = -0.21$ (with the male sex), $p_b = 0.04$ | 1 |
| Uniform Data Set (UDS)<br>*Number of Available Sessions* | L. Superior parietal cortex – L. Precuneus cortex<br>*Raw*, $r = 0.37$, $p_b = 5 \times 10^{-18}$ | 73 |
| | R. Inferior parietal cortex – L. Inferior parietal cortex<br>*Augmented*, $r = 0.34$, $p_b = 3 \times 10^{-13}$ | 1430 |
| Neuropsychological Assessment<br>*Number of Available Sessions* | L. Superior parietal cortex – L. Precuneus cortex<br>*Raw*, $r = 0.38$, $p_b = 9 \times 10^{-13}$ | 105 |
| | L. Pericalcarine cortex – L. Parahippocampal cortex<br>*Augmented*, $r = 0.41$, $p_b = 10^{-16}$ | 2519 |
| ADRC Clinical Data<br>*Number of Available Sessions* | L. Superior parietal cortex – L. Precuneus cortex<br>*Raw*, $r = 0.31$, $p_b = 4 \times 10^{-11}$ | 58 |
| | R. Inferior parietal cortex – R. Fusiform cortex<br>*Augmented*, $r = 0.31$, $p_b = 2 \times 10^{-11}$ | 1060 |
| Number of PET Sessions | R. Hippocampus – R. Fusiform cortex<br>*Raw*, $r = 0.3$, $p_b = 4 \times 10^{-7}$ | 19 |
| | L. Hippocampus – R. Cuneus cortex<br>*Augmented*, $r = 0.34$, $p_b = 7 \times 10^{-11}$ | 574 |
| Number of MRI Sessions | R. Hippocampus – R. Middle temporal cortex<br>*Raw*, $r = 0.35$, $p_b = 3 \times 10^{-15}$ | 74 |
| | R. Hippocampus – R. Precuneus cortex<br>*Augmented*, $r = 0.38$, $p_b = 10^{-20}$ | 1937 |
| Number of CT Sessions | R. Hippocampus – R. Superior frontal cortex<br>*Augmented*, $r = 0.25$, $p_b = 0.01$ | 1 |
| Presumed disease status at enrollment<br>(*higher is healthier*) | R. Hippocampus – R. Ventral diencephalon<br>*Raw*, $r = 0.25$, $p_b = 0.0004$ | 10 |
| | R. Hippocampus – L. Superior frontal cortex<br>*Augmented*, $r = 0.3$, $p_b = 7 \times 10^{-10}$ | 336 |
| Level of independence<br>(*higher is more dependent*) | R. Hippocampus – L. Hippocampus<br>*Augmented*, $r = -0.29$, $p_b = 4 \times 10^{-8}$ | 182 |
| Form A3 has been submitted previously<br>and there have been no changes. | R. Precuneus cortex – L. Precuneus cortex<br>*Raw*, $r = -0.31$, $p_b = 5 \times 10^{-7}$ | 39 |
| | R. Rostral anterior cingulate ctx – L. Temporal pole ctx<br>*Augmented*, $r = -0.37$, $p_b = 10^{-12}$ | 1822 |
| Mother living | L. Hippocampus – R. Superior frontal cortex<br>*Augmented*, $r = 0.25$, $p_b = 0.001$ | 8 |
| Father living | R. Thalamus – L. Isthmus cingulate cortex<br>*Raw*, $r = 0.26$, $p_b = 0.0007$ | 2 |
| Sibling 1 living | R. Hippocampus – R. Fusiform cortex<br>*Raw*, $r = 0.24$, $p_b = 0.02$ | 1 |
| | R. Hippocampus – R. Superior frontal cortex<br>*Augmented*, $r = 0.25$, $p_b = 0.006$ | 10 |
| Sibling 2 living | R. Hippocampus – R. Fusiform cortex<br>*Raw*, $r = 0.28$, $p_b = 0.04$ | 1 |
| Subject wearing a hearing aid | L. Lingual cortex – L. Putamen<br>*Raw*, $r = -0.21$, $p_b = 0.02$ | 1 |
| | R. Lingual cortex – L. Pallidum<br>*Augmented*, $r = -0.22$, $p_b = 0.006$ | 10 |
| History or presence of hypertension | R. Superior frontal cortex – R. Thalamus<br>*Augmented*, $r = -0.23$, $p_b = 0.001$ | 14 |
| Hachinski Ischemic score | R. Superior frontal cortex – R. Ventral diencephalon<br>*Raw*, $r = -0.22$, $p_b = 0.03$ | 1 |
| | R. Superior frontal cortex – R. Thalamus<br>*Augmented*, $r = -0.25$, $p_b = 9 \times 10^{-5}$ | 20 |
| Mini-Mental State Examination (MMSE) | R. Hippocampus – R. Putamen<br>*Raw*, $r = 0.29$, $p_b = 7 \times 10^{-9}$ | 40 |
| | R. Hippocampus – L. Precentral cortex<br>*Augmented*, $r = 0.38$, $p_b = 3 \times 10^{-20}$ | 702 |
| Clinical Dementia Rating (CDR)<br>*Memory* | R. Hippocampus – R. Amygdala<br>*Raw*, $r = -0.33$, $p_b = 3 \times 10^{-13}$ | 65 |
| | R. Hippocampus – L. Thalamus<br>*Augmented*, $r = -0.41$, $p_b = 7 \times 10^{-25}$ | 1499 |
| Clinical Dementia Rating (CDR)<br>*Orientation* | R. Hippocampus – R. Amygdala<br>*Raw*, $r = -0.3$, $p_b = 10^{-9}$ | 37 |
| | R. Hippocampus – L. Insula cortex<br>*Augmented*, $r = -0.37$, $p_b = 2 \times 10^{-18}$ | 1080 |
| Clinical Dementia Rating (CDR)<br>*Judgment and problem-solving* | R. Hippocampus – R. Amygdala<br>*Raw*, $r = -0.31$, $p_b = 2 \times 10^{-10}$ | 48 |
| | R. Hippocampus – R. Superior frontal cortex<br>*Augmented*, $r = -0.39$, $p_b = 3 \times 10^{-21}$ | 993 |
| Clinical Dementia Rating (CDR)<br>*Community affairs* | R. Hippocampus – R. Superior frontal cortex<br>*Augmented*, $r = -0.35$, $p_b = 2 \times 10^{-16}$ | 594 |
| Clinical Dementia Rating (CDR)<br>*Home and hobbies* | R. Hippocampus – R. Amygdala<br>*Raw*, $r = -0.29$, $p_b = 7 \times 10^{-9}$ | 38 |
| | R. Hippocampus – L. Thalamus<br>*Augmented*, $r = -0.38$, $p_b = 10^{-20}$ | 938 |
| Clinical Dementia Rating (CDR)<br>*Sum of boxes* | R. Hippocampus – R. Amygdala<br>*Raw*, $r = -0.32$, $p_b = 10^{-11}$ | 46 |
| | R. Hippocampus – L. Thalamus<br>*Augmented*, $r = -0.4$, $p_b = 2 \times 10^{-23}$ | 1199 |
| Clinical Dementia Rating (CDR)<br>*Total score* | R. Hippocampus – R. Amygdala<br>*Raw*, $r = -0.33$, $p_b = 10^{-12}$ | 55 |
| | R. Hippocampus – L. Thalamus<br>*Augmented*, $r = -0.41$, $p_b = 10^{-24}$ | 1163 |
| Difficulty or needing help with<br>*paying bills* | R. Hippocampus – R. Amygdala<br>*Raw*, $r = -0.27$, $p_b = 2 \times 10^{-6}$ | 19 |
| | R. Hippocampus – R. Superior frontal cortex<br>*Augmented*, $r = -0.33$, $p_b = 2 \times 10^{-12}$ | 579 |
| Difficulty or needing help with<br>*taxes and business affairs* | R. Hippocampus – R. Amygdala<br>*Raw*, $r = -0.27$, $p_b = 10^{-6}$ | 23 |
| | R. Hippocampus – L. Thalamus<br>*Augmented*, $r = -0.34$, $p_b = 5 \times 10^{-14}$ | 535 |
| Difficulty or needing help with<br>*shopping alone* | R. Hippocampus – R. Ventral diencephalon<br>*Raw*, $r = -0.25$, $p_b = 0.0004$ | 4 |
| | R. Hippocampus – L. Thalamus<br>*Augmented*, $r = -0.3$, $p_b = 10^{-9}$ | 225 |
| Difficulty or needing help with<br>*games and hobbies* | L. Hippocampus – L. Ventral diencephalon<br>*Raw*, $r = -0.22$, $p_b = 0.03$ | 1 |
| | L. Hippocampus – R. Hippocampus<br>*Augmented*, $r = -0.28$, $p_b = 2 \times 10^{-7}$ | 97 |
| Difficulty or needing help with<br>*preparing a balanced meal* | R. Hippocampus – R. Amygdala<br>*Raw*, $r = -0.23$, $p_b = 0.003$ | 4 |
| | R. Hippocampus – L. Pallidum<br>*Augmented*, $r = -0.3$, $p_b = 10^{-9}$ | 213 |
| Difficulty or needing help with<br>*keeping track of current events* | R. Hippocampus – R. Ventral diencephalon<br>*Raw*, $r = -0.23$, $p_b = 0.01$ | 1 |
| | R. Hippocampus – L. Thalamus<br>*Augmented*, $r = -0.28$, $p_b = 3 \times 10^{-8}$ | 112 |
| Difficulty or needing help with<br>*paying attention* | R. Hippocampus – R. Fusiform cortex<br>*Raw*, $r = -0.23$, $p_b = 0.002$ | 2 |
| | R. Hippocampus – L. Precuneus cortex<br>*Augmented*, $r = -0.29$, $p_b = 5 \times 10^{-9}$ | 369 |
| Difficulty or needing help with<br>*remembering dates* | R. Hippocampus – R. Ventral diencephalon<br>*Raw*, $r = -0.24$, $p_b = 0.001$ | 2 |
| | R. Hippocampus – L. Insula cortex<br>*Augmented*, $r = -0.29$, $p_b = 5 \times 10^{-9}$ | 106 |
| Difficulty or needing help with<br>*traveling and driving* | R. Hippocampus – R. Ventral diencephalon<br>*Raw*, $r = -0.29$, $p_b = 10^{-7}$ | 24 |
| | R. Hippocampus – R. Superior frontal cortex<br>*Augmented*, $r = -0.35$, $p_b = 6 \times 10^{-16}$ | 452 |
| Decline<br>*reported by subject* | L. Thalamus – L. Parahippocampal cortex<br>*Raw*, $r = -0.22$, $p_b = 0.02$ | 4 |
| | L. Hippocampus – R. Superior frontal cortex<br>*Augmented*, $r = -0.29$, $p_b = 6 \times 10^{-9}$ | 292 |
| Decline<br>*reported by informant* | R. Hippocampus – R. Amygdala<br>*Raw*, $r = -0.28$, $p_b = 10^{-7}$ | 40 |
| | L. Hippocampus – R. Superior frontal cortex<br>*Augmented*, $r = -0.39$, $p_b = 3 \times 10^{-20}$ | 1341 |
| Decline<br>*reported by clinician* | R. Hippocampus – R. Fusiform cortex<br>*Raw*, $r = -0.32$, $p_b = 10^{-10}$ | 66 |
| | R. Hippocampus – R. Superior frontal cortex<br>*Augmented*, $r = -0.43$, $p_b = 10^{-24}$ | 1611 |
| Cognitive impairment<br>*reported by clinician* | R. Hippocampus – R. Fusiform cortex<br>*Raw*, $r = -0.28$, $p_b = 0.0001$ | 18 |
| | L. Hippocampus – R. Lingual cortex<br>*Augmented*, $r = -0.36$, $p_b = 6 \times 10^{-11}$ | 223 |
| Wechsler Adult Intelligence Scale (WAIS)<br>*information* | R. Hippocampus – R. Inferior temporal cortex<br>*Raw*, $r = 0.26$, $p_b = 0.001$ | 9 |
| | R. Hippocampus – L. Caudate<br>*Augmented*, $r = 0.35$, $p_b = 10^{-11}$ | 344 |
| Wechsler Adult Intelligence Scale (WAIS)<br>*block design* | R. Hippocampus – R. Fusiform cortex<br>*Raw, $r = 0.36, p_b = 2 \times 10^{-12}$* | 54 |
|  | R. Hippocampus – L. Precentral cortex<br>*Augmented, $r = 0.4, p_b = 4 \times 10^{-17}$* | 1505 |
| Wechsler Adult Intelligence Scale (WAIS)<br>WAIS-R Digit Symbol | L. Hippocampus – L. Fusiform cortex<br>*Raw, $r = 0.39, p_b = 3 \times 10^{-15}$* | 96 |
|  | L. Hippocampus – R. Superior frontal cortex<br>*Augmented, $r = 0.45, p_b = 5 \times 10^{-24}$* | 1541 |
| Wechsler Memory Scale (WMS)<br>Associate Learning<br>*summary score* | L. Hippocampus – L. Thalamus<br>*Raw, $r = 0.31, p_b = 10^{-8}$* | 41 |
|  | L. Hippocampus – R. Superior frontal cortex<br>*Augmented, $r = 0.38, p_b = 3 \times 10^{-17}$* | 691 |
| Wechsler Memory Scale (WMS)<br>Digit Span Backward | L. Hippocampus – L. Thalamus<br>*Raw, $r = 0.25, p_b = 0.01$* | 1 |
|  | L. Hippocampus – L. Superior frontal cortex<br>*Augmented, $r = 0.27, p_b = 0.0005$* | 29 |
| Wechsler Memory Scale (WMS)<br>WMS-III Letter-Number Sequencing | L. Parahippocampal cortex – L. Ventral diencephalon<br>*Raw, $r = 0.26, p_b = 10^{-5}$* | 16 |
|  | R. Hippocampus – L. Superior frontal cortex<br>*Augmented, $r = 0.33, p_b = 10^{-12}$* | 255 |
| Total animals named in 60 seconds | R. Hippocampus – R. Ventral diencephalon<br>*Raw, $r = 0.27, p_b = 3 \times 10^{-6}$* | 29 |
|  | R. Hippocampus – L. Superior frontal cortex<br>*Augmented, $r = 0.35, p_b = 8 \times 10^{-16}$* | 350 |
| Total vegetables named in 60 seconds | L. Hippocampus – L. Inferior temporal cortex<br>*Raw, $r = 0.27, p_b = 5 \times 10^{-5}$* | 10 |
|  | R. Hippocampus – L. Precentral cortex<br>*Augmented, $r = 0.31, p_b = 6 \times 10^{-9}$* | 181 |
| Mental Control<br>*total score* | L. Hippocampus – R. Superior frontal cortex<br>*Augmented, $r = 0.27, p_b = 0.0001$* | 72 |
| Trail Making Test, Part A<br>*Time to complete* | R. Hippocampus – R. Fusiform cortex<br>*Raw, $r = -0.29, p_b = 3 \times 10^{-8}$* | 35 |
|  | R. Hippocampus – L. Superior frontal cortex<br>*Augmented, $r = -0.37, p_b = 5 \times 10^{-18}$* | 836 |
| Trail Making Test, Part B<br>*Time to complete* | L. Hippocampus – L. Fusiform cortex<br>*Raw, $r = -0.34, p_b = 2 \times 10^{-13}$* | 94 |
|  | L. Hippocampus – R. Superior frontal cortex<br>*Augmented, $r = -0.43, p_b = 3 \times 10^{-26}$* | 1643 |
| Boston Naming Test<br>*60 items* | R. Hippocampus – R. Inferior temporal cortex<br>*Raw, $r = 0.29, p_b = 2 \times 10^{-5}$* | 20 |
|  | R. Hippocampus – R. Thalamus<br>*Augmented, $r = 0.37, p_b = 2 \times 10^{-12}$* | 1142 |
| Current Logical Memory IA<br>*story units recalled* | R. Hippocampus – R. Amygdala<br>*Raw, $r = 0.3, p_b = 7 \times 10^{-6}$* | 13 |
|  | R. Hippocampus – L. Ventral diencephalon<br>*Augmented, $r = 0.36, p_b = 4 \times 10^{-11}$* | 478 |
| Logical Memory IIA – Delayed<br>*story units recalled* | R. Hippocampus – R. Amygdala<br>*Raw, $r = 0.31, p_b = 5 \times 10^{-7}$* | 26 |
|  | R. Hippocampus – L. Ventral diencephalon<br>*Augmented, $r = 0.38, p_b = 3 \times 10^{-13}$* | 757 |
| Simon<br>*percent correct* | R. Hippocampus – L. Hippocampus<br>*Augmented, $r = 0.25, p_b = 0.0003$* | 36 |
| Simon<br>*number of correct on all trials* | L. Hippocampus – R. Superior frontal cortex<br>*Augmented, $r = 0.26, p_b = 4 \times 10^{-5}$* | 53 |
| Switch pure CV<br>*number correct* | L. Hippocampus – R. Superior parietal cortex<br>*Augmented, $r = 0.23, p_b = 0.006$* | 7 |
| Switch mixed<br>*number correct* | L. Hippocampus – L. Ventral diencephalon<br>*Raw, $r = 0.26, p_b = 0.0002$* | 15 |
|  | L. Hippocampus – R. Caudate<br>*Augmented, $r = 0.34, p_b = 2 \times 10^{-11}$* | 205 |
| Switch<br>*percent correct* | L. Hippocampus – L. Fusiform cortex<br>*Raw, $r = 0.27, p_b = 2 \times 10^{-5}$* | 11 |
|  | L. Hippocampus – R. Precuneus cortex<br>*Augmented, $r = 0.34, p_b = 8 \times 10^{-13}$* | 338 |
| Benson Complex Figure Copy | R. Hippocampus – L. Inferior temporal cortex<br>*Augmented, $r = 0.26, p_b = 0.02$* | 3 |
| Craft Story 21 Recall (Immediate)<br>*verbatim scoring* | R. Hippocampus – R. Superior frontal cortex<br>*Augmented, $r = 0.29, p_b = 0.0008$* | 6 |
| Craft Story 21 Recall (Immediate)<br>*paraphrase scoring* | R. Hippocampus – R. Superior frontal cortex<br>*Augmented, $r = 0.32, p_b = 9 \times 10^{-6}$* | 42 |
| Craft Story 21 Recall (Delayed)<br>*verbatim scoring* | R. Hippocampus – R. Superior frontal cortex<br>*Augmented, $r = 0.28, p_b = 0.003$* | 8 |
| Craft Story 21 Recall (Delayed)<br>*paraphrase scoring* | R. Hippocampus – R. Superior frontal cortex<br>*Augmented, $r = 0.3, p_b = 0.0002$* | 26 |
| Multilingual Naming Test (MINT)<br>*total score* | R. Hippocampus – L. Caudate<br>*Augmented, $r = 0.28, p_b = 0.04$* | 1 |
| Multilingual Naming Test (MINT)<br>*total correct without semantic cue* | R. Hippocampus – L. Caudate<br>*Augmented, $r = 0.28, p_b = 0.04$* | 1 |
| Multilingual Naming Test (MINT)<br>*phonemic cues: number given* | R. Hippocampus – L. Caudate<br>*Augmented, $r = -0.3, p_b = 0.004$* | 4 |
| Montreal Cognitive Assessment (MoCA)<br>*Total Raw Score – uncorrected* | R. Hippocampus – R. Thalamus<br>*Raw, $r = 0.31, p_b = 4 \times 10^{-5}$* | 8 |
|  | R. Hippocampus – R. Superior frontal cortex<br>*Augmented, $r = 0.32, p_b = 3 \times 10^{-7}$* | 95 |
| Montreal Cognitive Assessment (MoCA)<br>*Delayed recall – no cue* | R. Hippocampus – R. Thalamus<br>*Raw, $r = 0.31, p_b = 3 \times 10^{-5}$* | 3 |
|  | R. Hippocampus – L. Superior frontal cortex<br>*Augmented, $r = 0.26, p_b = 0.006$* | 4 |
| Free and Cued Selective Reminding Test<br>*Trial 1 Free Recall* | R. Hippocampus – R. Precentral cortex<br>*Raw, $r = 0.33, p_b = 8 \times 10^{-12}$* | 30 |
|  | R. Hippocampus – L. Thalamus<br>*Augmented, $r = 0.34, p_b = 2 \times 10^{-13}$* | 158 |
| Free and Cued Selective Reminding Test<br>*Trial 1 Cued Recall* | R. Hippocampus – R. Precentral cortex<br>*Raw, $r = -0.25, p_b = 0.0002$* | 2 |
|  | R. Hippocampus – R. Precentral cortex<br>*Augmented, $r = -0.21, p_b = 0.03$* | 3 |
| Free and Cued Selective Reminding Test<br>*Trial 2 Free Recall* | L. Hippocampus – L. Ventral diencephalon<br>*Raw, $r = 0.32, p_b = 4 \times 10^{-10}$* | 38 |
|  | R. Hippocampus – L. Hippocampus<br>*Augmented, $r = 0.38, p_b = 10^{-18}$* | 350 |
| Free and Cued Selective Reminding Test<br>*Trial 2 Cued Recall* | R. Hippocampus – R. Precentral cortex<br>*Raw, $r = -0.28, p_b = 2 \times 10^{-6}$* | 16 |
|  | L. Hippocampus – R. Superior frontal cortex<br>*Augmented, $r = -0.3, p_b = 4 \times 10^{-10}$* | 94 |
| Free and Cued Selective Reminding Test<br>*Trial 3 Free Recall* | R. Hippocampus – R. Putamen<br>*Raw, $r = 0.31, p_b = 7 \times 10^{-10}$* | 41 |
|  | R. Hippocampus – R. Superior frontal cortex<br>*Augmented, $r = 0.39, p_b = 5 \times 10^{-21}$* | 687 |
| Free and Cued Selective Reminding Test<br>*Trial 3 Cued Recall* | R. Hippocampus – R. Precentral cortex<br>*Raw, $r = -0.28, p_b = 10^{-6}$* | 26 |
|  | R. Hippocampus – R. Superior frontal cortex<br>*Augmented, $r = -0.34, p_b = 6 \times 10^{-14}$* | 233 |
| Free and Cued Selective Reminding Test<br>*free summary score* | R. Hippocampus – R. Precentral cortex<br>*Raw, $r = 0.35, p_b = 4 \times 10^{-13}$* | 44 |
|  | R. Hippocampus – R. Superior frontal cortex<br>*Augmented, $r = 0.39, p_b = 7 \times 10^{-21}$* | 541 |
| Free and Cued Selective Reminding Test<br>*total score* | R. Hippocampus – R. Ventral diencephalon<br>*Raw, $r = 0.23, p_b = 0.02$* | 2 |
|  | R. Hippocampus – L. Thalamus<br>*Augmented, $r = 0.27, p_b = 10^{-6}$* | 167 |
| Clinician Diagnosis:<br>Normal Cognition | R. Hippocampus – R. Amygdala<br>*Raw, $r = 0.32, p_b = 2 \times 10^{-11}$* | 69 |
|  | R. Hippocampus – R. Superior frontal cortex<br>*Augmented, $r = 0.41, p_b = 3 \times 10^{-24}$* | 1648 |
| Clinician Diagnosis:<br>Alzheimer's Disease | R. Hippocampus – R. Fusiform cortex<br>*Raw, $r = -0.3, p_b = 6 \times 10^{-6}$* | 19 |
|  | L. Hippocampus – R. Lingual cortex<br>*Augmented, $r = -0.35, p_b = 2 \times 10^{-10}$* | 308 |
| Amyloid burden (Centiloid scale)<br>*mean cortical*<br>*binding potential* | L. Parahippocampal cortex – L. Ventral diencephalon<br>*Raw, $r = -0.27, p_b = 0.007$* | 2 |
|  | L. Hippocampus – R. Superior temporal cortex<br>*Augmented, $r = -0.31, p_b = 2 \times 10^{-5}$* | 46 |
| Amyloid burden (Centiloid scale)<br>*mean cortical*<br>*standardized uptake value ratio (SUVR)* | L. Parahippocampal cortex – L. Ventral diencephalon<br>*Raw, $r = -0.26, p_b = 0.0003$* | 8 |
|  | L. Hippocampus – R. Superior temporal cortex<br>*Augmented, $r = -0.31, p_b = 10^{-7}$* | 144 |
| Amyloid burden (Centiloid scale)<br>*mean cortical*<br>*binding potential*<br>*(partial-volume corrected)* | L. Parahippocampal cortex – L. Ventral diencephalon<br>*Raw, $r = -0.28, p_b = 0.001$* | 4 |
|  | L. Hippocampus – R. Superior temporal cortex<br>*Augmented, $r = -0.34, p_b = 10^{-7}$* | 86 |
| Amyloid burden (Centiloid scale)<br>*mean cortical*<br>*standardized uptake value ratio (SUVR)*<br>*(partial-volume corrected)* | L. Parahippocampal cortex – L. Ventral diencephalon<br>*Raw, $r = -0.29, p_b = 5 \times 10^{-6}$* | 20 |
|  | L. Hippocampus – R. Superior temporal cortex<br>*Augmented, $r = -0.35, p_b = 6 \times 10^{-12}$* | 608 |

**Table 3:** Significant correlations of non-MRI variables with brain connectivity in PREVENT-AD.

| Non-MRI variable | Most correlated brain structural connection | # Sig. Conn |
| --- | --- | --- |
| Age | R. Hippocampus – R. Thalamus<br>*Raw*, $r = -0.33$, $p_b = 0.002$ | 2 |
| | L. Hippocampus – R. Thalamus<br>*Augmented*, $r = -0.33$, $p_b = 0.001$ | 13 |
| Age of mother at AD-like dementia onset | R. Ventral diencephalon – L. Banks of superior temporal sulcus<br>*Raw*, $r = -0.36$, $p_b = 0.006$ | 1 |
| Tau phosphorylated at Thr181 (P-tau) concentration in CSF | R. Caudate – L. Caudal middle frontal cortex<br>*Raw*, $r = 0.45$, $p_b = 0.04$ | 1 |

**Table 4:** Significant correlations of non-MRI variables with brain connectivity in HCP.

| Non-MRI variable | Most correlated brain structural connection | # Sig. Conn |
| --- | --- | --- |
| Sex | Brainstem – L. Paracentral cortex<br>*Raw*, $r = -0.24$ (with the male sex), $p_b = 0.007$ | 2 |
| Height | R. Parahippocampal cortex – L. Paracentral cortex<br>*Augmented*, $r = -0.22$, $p_b = 0.0497$ | 1 |
| Weight | R. Ventral diencephalon – L. Ventral diencephalon<br>*Raw*, $r = -0.24$, $p_b = 0.008$ | 5 |

- initially not found in ADNI-2 but appeared as positive by including ICV as a covariate,
- negative in OASIS-3 regardless of controlling for ICV (but stronger without),
- initially positive in PREVENT-AD but disappeared after including ICV as a covariate,
- negative in HCP regardless of controlling for ICV (but stronger without).

Next, we focused on three representative variables of age, MMSE, and CDR (sum of boxes). Among all the augmented brain connections that were significantly correlated with age (after Bonferroni correction) consistently in ADNI-2, OASIS-3, and PREVENT-AD, the one between the right superior frontal cortex and the left hippocampus was most significant (in terms of the geometric mean of the *p*-values across databases). Similarly, for both MMSE and CDR, the augmented connection between the right superior frontal cortex and the right hippocampus had the most significant correlation consistently in ADNI-2 and OASIS-3. These relationships are plotted in Figure 2 for all three variables.

**Figure 2.**
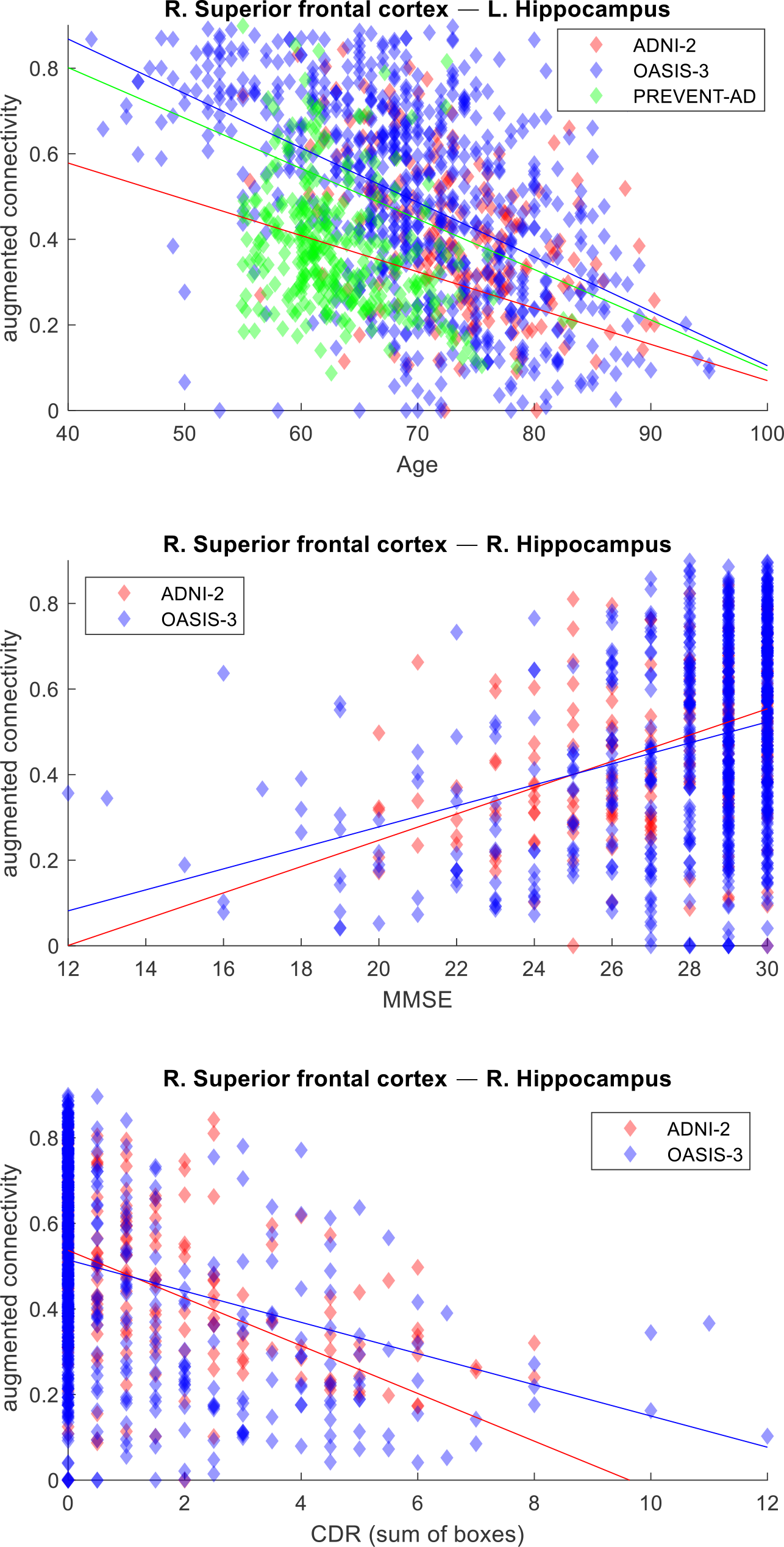
Augmented connectivity of the brain connection most significantly correlated with age (top), MMSE (middle), and CDR (bottom) consistently across databases.

## 4. Discussion

Although more correlations were initially found to be significant with raw than augmented structural connectivity (in three out of four databases), visual inspection of the data led to discarding many of the former – but none of the latter – as spurious, implying more robustness and reliability of the augmented structural connections. Spuriousness was often because raw (direct) connectivity between an ROI pair was zero for all except a few subjects that dramatically influenced the correlation calculation, in contrast to augmented connectivity that is always positive in a network with a single connected component. Eventually, a total of 85 relationships with raw connectivity and 101 with augmented connectivity passed the Pearson correlation threshold, Bonferroni correction, and visual inspection. Each variable may have been significantly correlated with multiple brain connections, the number of which is listed in the tables (right column). Out of 76 variables correlated with both types of connectivity (see taller cells in the left column of the tables), 72 were more significantly correlated with augmented than raw connectivity, always with a greater or equal number of total connections correlated with augmented than raw connectivity. For a significance plot of the correlation with age, MMSE, and CDR for all connections (albeit in a non-exploratory context and with a different connectivity quantification method), see our previous report [18].

More variables were found to be significantly related to brain connectivity in ADNI-2 and OASIS-3 than in PREVENT-AD and HCP, possibly due to the fact that PREVENT-AD and HCP (that include only healthy subjects) are more homogenous populations with narrower ranges of scores (e.g. MMSE) than ADNI-2 and OASIS-3 (that include a mix of healthy, MCI, and AD subjects). Moreover, the fact that proxies for disease severity, such as MMSE and CDR, correlate with brain connectivity (only with the presence of MCI and AD patients) suggests that the corresponding changes in the connectome are possibly disease-related and a potential marker of the disease.

The most prominent non-MRI variable that was consistently correlated with structural connectivity was age. A negative correlation was observed between age and hippocampal connectivity in all databases except HCP. The limited age range in the young population of HCP may be the reason why this relationship was not detected in this database, given that the standard deviations of age were (in decreasing order) 9.1 years in OASIS-3, 6.9 years in ADNI-2, 5.1 years in PREVENT-AD, but only 3.6 years in HCP. In fact, the statistical significance of the age correlation decreased in the same database order.

Clinical scores that were found to be significantly related to brain connectivity showed the consistent trend of healthier score being linked to increased connectivity. The only exception was the significant relationship of P-tau with the raw connection between the right caudate and the left caudal middle frontal cortex in PREVENT-AD, which was unanticipatedly *positive*. Nonetheless, we had already observed – in a different database with a different connectivity quantification method – a similarly unexpected strengthening of caudal structural connectivity with worsening cognitive status [18, 37]. In fact, volume [38] and fractional anisotropy (FA) [39] of the caudate have been reported to increase in pre-symptomatic familial AD, which might have also led to the aforementioned relationship we observed in PREVENT-AD (that includes healthy subjects at risk of AD). Such an increase in the measured structural connectivity in pre-symptomatic subjects may indicate a compensatory effect [40], or could stem from other factors (e.g., selective axonal loss can increase FA in regions with fiber crossing [39, 41, 42]).

The number of imaging sessions and clinical data available for a subject in OASIS-3 was positively related to (mostly) hippocampal connectivity. This could be attributable to a higher follow-up rate for those with healthier hippocampi, as individuals with MCI and dementia have been shown to have lower retention rates in research studies than those with normal cognition [43, 44].

With larger ICV, brain regions become farther apart from each other, thus harder to reach by streamline tractography. Therefore, we decided to control for ICV in our regression analysis to avoid underestimation of brain connectivity. Doing so eliminated (in some databases) correlation of brain connectivity with several variables, some of which might have been spurious due to possible correlation with ICV, e.g., strength, sex, alcohol consumption, and posture. Correlation of brain connectivity with sex [45, 46], in particular, remained inconclusive, given that it disappeared in PREVENT-AD (and was weakened in OASIS-3 and HCP) after ICV adjustment, as is typically seen in neuroimaging studies [47, 48], and appeared in ADNI-2 only after ICV adjustment, which could be a sign of an introduced (previously absent) ICV bias [49] (especially as the direction of the relationship in ADNI-2 was opposite to that in OASIS-3).

Differences in scanner hardware, population characteristics, and protocols across databases introduce database-dependent effects on both the acquired images and the measured variables (see the figures). Such effects could create large variances in both measured brain connectivity and non-MRI variables (those common in multiple databases) if databases were combined in a single heterogeneous correlation analysis. To prevent the natural population-level variances – that lead to true correlations – from being overshadowed by heterogeneity variances due to multi-database combination, we decided to analyze the databases independently and then compare the findings. Although data harmonization [50] in a combined database setting could reduce the heterogeneity to some extent, it would take away the benefit of replicability assessment across databases. Note that even within a single database, data may come from various sites; however, the within-database site effect is expected to be smaller, as all sites supposedly often follow the same database-wide protocol. Data harmonization could nevertheless increase the statistical power when analyzing a multi-site database.

To correct for multiple comparisons, we took the conservative Bonferroni approach to avoid false-positive relationships with brain connectivity (reduce type I errors). Strong interdependencies both between brain connections and between non-MRI variables can nevertheless be exploited to design a complex but more forgiving correction scheme in order to avoid false negatives and missing existing relationships (reduce type II errors).

## 5. Conclusions

We conducted a retrospective exploratory study to examine the associations between brain structural connectivity and non-MRI variables, using data from four (including three AD-related) public dMRI databases. Unlike hypothesis-driven research, where conjectured relationships between specific variables are tested, we calculated the correlation between all brain connections and non-MRI variables in our dataset without prior assumption, while stringently correcting for multiple comparisons, with the aim of discovering connectomic relationships. Replication of our findings in other databases (such as ADNI-3) and with other connectivity quantification methods and conducting our study with harmonized dMRI data are subjects of future research.

## Acknowledgements

Support for this research was provided by the National Institutes of Health (NIH), specifically the National Institute on Aging (NIA; R56AG068261, RF1AG068261).

Additional support was provided in part by the BRAIN Initiative Cell Census/Atlas Network grants U01MH117023 and UM1MH130981, the National Institute for Biomedical Imaging and Bioengineering (NIBIB; P41EB015896, R01EB023281, R01EB006758, R21EB018907, R01EB019956, P41EB030006), the NIA (R21AG082082, R56AG064027, R01AG064027, R01AG008122, R01AG016495, R01AG070988), the National Institute of Mental Health (R01MH121885, RF1MH123195), the National Institute for Neurological Disorders and Stroke (R01NS0525851, R21NS072652, R01NS070963, R01NS083534, U01NS086625, U24NS10059103, R01NS105820), the NIH Blueprint for Neuroscience Research (U01MH093765), part of the multi-institutional Human Connectome Project, the BrightFocus Foundation (A2016172S), and the Michael J. Fox Foundation for Parkinson’s Research (MJFF-021226).

Computational resources were provided through the Massachusetts Life Sciences Center and NIH Shared Instrumentation Grants (S10RR023401, S10RR019307, S10RR023043, S10RR028832).

Data collection and sharing for this project was funded in part by the Alzheimer’s Disease Neuroimaging Initiative (ADNI) (NIH Grant U01 AG024904) and DOD ADNI (Department of Defense award number W81XWH-12-2-0012). ADNI is funded by the NIA, the NIBIB, and through generous contributions from the following: AbbVie, Alzheimer’s Association; Alzheimer’s Drug Discovery Foundation; Araclon Biotech; BioClinica, Inc.; Biogen; Bristol-Myers Squibb Company; CereSpir, Inc.; Cogstate; Eisai Inc.; Elan Pharmaceuticals, Inc.; Eli Lilly and Company; EuroImmun; F. Hoffmann-La Roche Ltd and its affiliated company Genentech, Inc.; Fujirebio; GE Healthcare; IXICO Ltd.; Janssen Alzheimer Immunotherapy Research & Development, LLC.; Johnson & Johnson Pharmaceutical Research & Development LLC.; Lumosity; Lundbeck; Merck & Co., Inc.; Meso Scale Diagnostics, LLC.; NeuroRx Research; Neurotrack Technologies; Novartis Pharmaceuticals Corporation; Pfizer Inc.; Piramal Imaging; Servier; Takeda Pharmaceutical Company; and Transition Therapeutics. The Canadian Institutes of Health Research is providing funds to support ADNI clinical sites in Canada. Private sector contributions are facilitated by the Foundation for the National Institutes of Health (www.fnih.org). The grantee organization is the Northern California Institute for Research and Education, and the study is coordinated by the Alzheimer’s Therapeutic Research Institute at the University of Southern California. ADNI data are disseminated by the Laboratory for Neuro Imaging at the University of Southern California.

Data were also provided in part by OASIS-3: Longitudinal Multimodal Neuroimaging: Principal Investigators: T. Benzinger, D. Marcus, J. Morris; NIH P30 AG066444, P50 AG00561, P30 NS09857781, P01 AG026276, P01 AG003991, R01 AG043434, UL1 TR000448, R01 EB009352.

AV-45 doses were provided by Avid Radiopharmaceuticals, a wholly owned subsidiary of Eli Lilly.

Data were also obtained in part from the Pre-symptomatic Evaluation of Novel or Experimental Treatments for Alzheimer’s Disease (PREVENT-AD) program.

Data were furthermore provided in part by the Human Connectome Project (HCP), WU-Minn Consortium (Principal Investigators: David Van Essen and Kamil Ugurbil; 1U54MH091657) funded by the 16 NIH Institutes and Centers that support the NIH Blueprint for Neuroscience Research; and by the McDonnell Center for Systems Neuroscience at Washington University.

B. Fischl has a financial interest in CorticoMetrics, a company whose medical pursuits focus on brain imaging and measurement technologies. His interests were reviewed and are managed by Massachusetts General Hospital and Mass General Brigham in accordance with their conflict-of-interest policies.

